# Development of a marker within the candidate *Un8* true loose smut resistance gene for use in barley breeding

**DOI:** 10.1101/2023.04.06.535815

**Authors:** Vilnis Šķipars, Elīna Sokolova, Sanita Seile, Dainis Ruņģis, Linda Legzdiņa

## Abstract

Breeding for resistance to true loose smut infection caused by the pathogen (*Ustilago nuda* (Jens.) Rostr.) is an economical and environmentally safe way to limit the effect of this pathogen on barley. However, screening for resistance using natural infection can lead to inconsistent results and artificial inoculation is labour intensive, and can fail, leading to erroneous phenotyping. Marker assisted selection of genes conferring disease resistance can increase the efficiency of breeding programs. A candidate gene for *Un8* resistance was used to develop a genetic marker which was tested on a recombinant inbred line (RIL) population derived from the resistant ‘CDC Freedom’ and the susceptible ‘Samson’ varieties. The RIL population (98 lines) was phenotyped for resistance to true loose smut by artificial inoculation and genotyped with the newly developed marker. The genotyping results obtained with the marker developed in this study were mostly consistent with true loose smut resistance determined by artificial inoculation. The markers was also tested in additional barley cultivars and breeding material. Repeated analysis of inconsistent results is required to confirm or revise these results, as well as further investigation of the candidate gene to confirm its role in barley true loose smut resistance is being planned.

## Introduction

Barley was domesticated approximately 10.000 years ago (Andersen et al., 2016) and now it is the fourth most important crop in the world (Nadolska-Orczyk1 et al., 2017, Asaad et al., 2013). Barley is one of the most adaptable cereal crops (Zhou, 2010), and is cultivated over a wide range of latitudes (Haas et al., 2019, Gürel et al., 2016). In recent years, barley has gained a renewed interest as a health food (Zhou, 2010; Day, 2013; Izydorczyk et al., 2019). Whole grain barley is highly nutritious and contains high levels of biological active compounds including β-glucan, phytochemicals, flavonoids, tochopherols, dietary fibers, tuocols (Sharma, 2019; Idehen et al., 2017). It is also increasingly studied and used for biofuel/ bioethanol production (Nghiem, 2010; Panahi, 2020; Lara-Serrano, 2018). Barley also has good potential to be used as an annual plant for paper production in straw-based cellulose factories (Munck, 1993).

The basidiomycete pathogen *Ustilago nuda* (Jens.) Rostr. (*U*.*nuda*) infects barley in all cultivation areas worldwide (Eckstein et al., 2002; Asaad et al., 2013). In epidemic years, *U*.*nuda* has caused yield losses of more than 30% (Asaad et al., 2013). Currently most grown cultivars are susceptible to loose smut. Disease controlling is done by mainly using uninfected seed or seed coated with systematic fungicides (Asaad et al., 2013; Eckstein et al., 2002). If infected seed is treated with fungicide, it reduces crop quality and quantity. Besides, continued fungicide use can lead to the rise of fungicide resistant mutants (Eckstein et al., 2002). In recent years, organic farming has become highly important and that has renewed an interest in seed-borne disease resistant crops (Wunderle et al., 2012). Breeding for loose smut resistance in barley for organic farming is particularly needed, as no seed treatments certified for organic farming are currently available (Muller, 2006).

*U*.*nuda* colonizes florets and as a dormant mycelium can overwinter in mature seeds (Zang et al., 2015). Upon seed germination, the mycelium aggregates into the florets – all parts of the ear, except for the rachis, are replaced by spores which are produced by segmentation of the hyphae (Malik and Bats, 1960). Plant remains symptomless until heading when the spikelets of the inflorescence are entirely replaced by a distinctive brown-black teliospore mass (Zang et al., 2015; Eckestein et al., 2002). The teliospores mature at about the same time when barley head emerges and are easily spread by the wind to other developing barley seeds where they infect healthy embryos and remain dormant until the seed germinates (Eckstein et al., 2002; Wunderle et al., 2012). Infection effectively takes place immediately before pollination until 4-8 days after (Pedersen, 1960; Wunderle et al., 2012). Infection rates are increased in cool high-rainfall areas and the year following a wet spring (Asaad et al., 2013).

As barley loose smut it is seed-born disease, cultivation of a pathogen free seed is the best method for reducing its spread (Asaad et al., 2013) and breeding for loose smut resistance is urgently needed because of the growing market for organic agriculture products (Muller, 2006). In addition, resistant varieties may lose their resistance to *U*.*nuda* by the appearance of new *U*.*nuda* pathotypes, as occurred with the Canadian barley variety ‘Titan’ (Skoropad et al., 1952).

DNA markers can be used to detect the presence of allelic variations in the genes underlying traits (Collard and Mackill, 2007). Molecular marker assisted selection (MAS) for resistance traits can greatly increase the efficiency and accuracy of breeding programmes, as natural inoculation is inconsistent and artificial inoculation is very time and labour consuming, requires two growing seasons and can fail due to poor germination of inoculated seeds. Therefore, the use of MAS for true loose smut resistance has the potential to be simpler than phenotypic screening and assessment of resistance and selection can be carried out at the seedling stage in off-season nurseries, enabling the growth of multiple generations per year (Zang et al., 2015; Collard and Mackill, 2007).

At least 15 genes controlling race specific resistance to *U*.*nuda* have been identified (Zang et al., 2015; Legkun et al., 2016). Most display dominant inheritance, with the exception of *un7*, which is recessive (Legkun et al., 2016). *Un8* has been reported to confer resistance to all known isolates of *U*.*nuda*, and is considered to be one of the most effective loose smut resistance genes (Eckstein et al., 2002; Zang et al., 2015; Legkun et al., 2016). A marker previously mapped to the *Un8* gene (Eckstein et al. 2002) has been shown to incompletely segregate with *Un8* conferred resistance, and is located several centimorgans (cM) from the gene, and is thus often not in agreement with the data from natural and artificial inoculation. A *Un8* candidate gene was identified in a 50kB interval on chromosome arm 1HL (Zang et al., 2015). This candidate gene encodes a tandem kinase protein, which have recently been revealed to be an important class of proteins that are involved in plant resistance (Klymiuk et al. 2021). A marker that is tightly linked to *Un8* co-segregates perfectly with *Un8* in a single mapping population, but this candidate gene has not been definitively proved to be the *Un8* resistance gene (Zang et al. 2015; Klymiuk et al. 2021).

The aim of this study was to develop a marker tightly linked with the *Un8* resistance locus in Latvian breeding germplasm, which can be used at the seedling stage and in the greenhouse several times per year. We developed a marker from the candidate *Un8* gene, and tested co-segregation with loose smut resistance in a recombinant inbred line (RIL) barley population created in Latvia, as well as additional varieties and breeding lines derived from the Latvian barley breeding program.

## Materials and Methods

### Recombinant Inbred Line (RIL) development

The RILs were developed using the resistant parent ‘CDC Freedom’ that has been verified to be resistant to true loose smut according to the information provided by the breeder and artificial inoculation (Muller, 2006), and ‘Samson’ was chosen as the susceptible parent (Table 1). After the initial cross the population was advanced from the F_1_ to F_5_ generations using the pedigree breeding method. A single seed or a single spike from each plant was harvested and sown to obtain the next generation from which again a single seed/spike from each headrow was harvested (F_2_ – F_5_). Starting from the F_6_ generation, each line was harvested and resown in bulk. From the 150 F_5_ RIL lines, 98 survived until the F_10_ generation, and these, together with both parental cultivars were used for genotyping. Artificial inoculation for true loose smut resistance was done in the F_8_ generation.

**Table 1.** Characteristics of the parental cultivars used to develop the RILs.

| **RIL parental cultivars** | **‘CDC Freedom’** | **‘Samson’** |
| --- | --- | --- |
| Country of origin | Canada | USA |
| True loose smut reaction (1) | R | S |
| *Un8* candidate gene allele group (2) | I | II |
| Row type | 2 | 6 |
| Grain type | Hulless | Hulled |
| Susceptibility to sprouting | S | R |

(1) Phenotypic reaction to true loose smut evaluated by artificial inoculation as described by Muller et al., (2006); (2) The Group I allele is present in barley lines resistant to true loose smut while the Group II–IV alleles are present in barley lines susceptible to true loose smut.

### Artificial inoculation

Artificial inoculations for assessment of resistance to true loose smut for 98 RILs were performed at Priekuli, Latvia in 2013 and 2014 as described by Mueller (2006). Additional lines known to be susceptible to the pathogen were tested as controls to monitor the inoculations. Inoculation was done on the field immediately prior to anthesis with a previously collected local mixture of true loose smut pathotypes of *U*.*nuda*. Inoculum was prepared using 1 g of teliospores per litre of distilled water (1g 1L ^-1^) and at least 4 ears per sample were pricked using a 10 ml syringe until the suspension emerged from the top of the floret. The inoculated plants were tagged and grown to maturity. The resistance phenotypes were scored in the next year during heading time. Artificial inoculation and spikes of individuals susceptible to barley loose smut are shown in Figure 1.

**Fig. 1.**
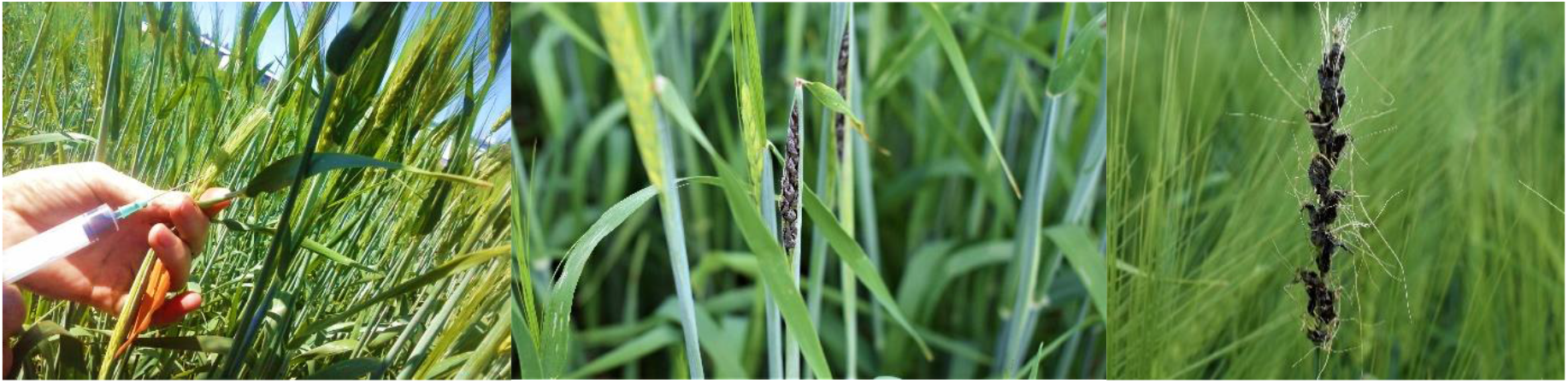
Artificial inoculation and spikes of individuals susceptible to barley loose smut.

### DNA extraction

Leaf samples were collected before flowering, five leaves per plant, and frozen immediately at - 20 °C. DNA extraction was done using a modified Wulff *et al*., (2002) method with the following changes: cleaning step with chloroform and isoamylalcohol was done once; ammonium acetate was omitted from washing buffer II; TE buffer was used only for storage of DNA. DNA quality and concentration was evaluated spectrophotometrically with a NanoDrop 2000 (ThermoFisher Scientific). Samples were re-extracted if DNA concentration was low or optic density ratio 260/280 nm was not in the optimal range of 1.8 – 2.1.

### Amplification and sequencing of candidate gene for *Un8* gene

The *Un8* candidate gene was Sanger sequenced from both parents of the RIL using the PCR primer pairs described in Zang et al., (2015) (0751D06_1 and 0751D06_2). These primer pairs span the entire coding region of the *Un8* candidate gene as well as portions of the 5’ and 3’ untranslated regions, primer pair 0751D06_1, amplifying the region from −292 to +1039, and primer pair 0751D06_2 amplifying the region from +957 to +2103. Positive numbers refer to the nucleotide position relative to the adenosine nucleotide of the translation start codon (position 1). Position numbers are according to the reference gene sequence in which the coding region was 2037 nucleotides in length.

PCR amplification was done in a total volume of 10 μl containing 2 μl of 5x HOT FIREPol® Blend Master Mix Ready to Load (Solis BioDyne, Estonia), MgCl_2_ (final concentration 2 mM), PCR primers with final concentration of 200 nM. The PCR amplification profile was: 95 °C for 15’ followed by 35 cycles of 95 °C for 45 s, 64 °C for 45 s and 72 °C for 1 min, followed by a final elongation step of 5 min at 72 °C. Prior to Sanger sequencing, PCR products were treated with ExoSAP-IT™ Express PCR Product Cleanup reagent (Thermo Fisher Scientific Baltics, Lithuania), according to the manufacturer’s protocol. The Sanger sequencing reactions were carried out using the BrilliantDye™ Terminator (v1.1) Cycle Sequencing kit (NimaGen, the Netherlands), following the manufacturer’s instructions and analysed using the Genetic Analyzer 3500 (Thermo Fisher Scientific, USA).

Sequences from the RIL parental varieties were aligned with the reference *Un8* sequence MLOC_38442 (Zang et al, 2015). ContigExpress and AlignX modules from the VectorNTI software (Thermo Fisher Scientific, USA) were used for sequence alignment. The consensus sequences obtained from the resistant parental cultivar (‘CDC Freedom’), and the susceptible parental cultivar (‘Samson’), were deposited in the National Center for Biotechnology Information (NCBI) GenBank nucleic acid sequence database (accession nos. TBC).

### Marker development and genotyping

Based on nucleotide sequence differences between the susceptible and resistant parental varieties of the tested RIL population, a restriction enzyme based genotyping assay was developed. A 170 bp long PCR product was created using the primers *Un8*_PvuII-F (5’-GTACATGCACGAGGAGTGCT-3’) and *Un8*_PvuII-R (5’-ACTCAGGCGCCATATACCC-3’).

Total PCR volume was 10 μl: each primer with final concentration 250 nM, 2 μl of 5x HOT FIREPol® Blend Master Mix (Solis BioDyne, Estonia), MgCl_2_ with final concentration of 2 mM, 1 μl of DNA sample of concentration (11.5 – 18.5 ng / μl). PCR cycling conditions were: 95 °C 15’ followed by 35 cycles of 95 °C 10 s, 62 °C 20 s and 72 °C 30 s. The amplified PCR fragments were digested with the PvuII restriction enzyme (Thermo Fisher Scientific Baltics, Lithuania) and separated on a 2% agarose gel (in 1X TAE buffer), stained with EtBr. To perform the restriction, 22 μl of reaction mixture (18 μl of deionized water, 2 μl of 10X buffer G and 20 U of PvuII per sample) were added to the PCR products followed by incubation at 37 °C for 2 h.

The RIL population was genotyped using the developed assay. Individuals with 111bp and 59bp digestion fragments were considered as homozygous resistant and individuals with only the 170bp fragment as homozygous susceptible. Individuals where all three fragments (59; 111; 170 bp) were observed were regarded as heterozygous (Fig. 2).

**Fig. 2.**
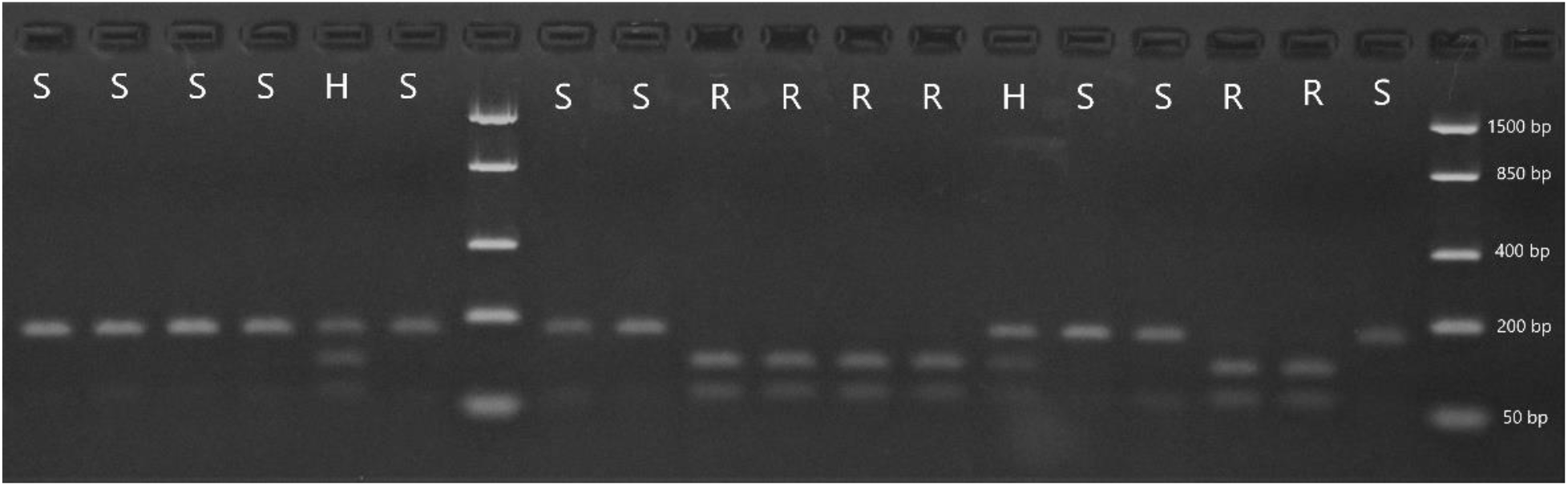
Cleavage of PCR products from RIL population with PvuII. R – resistant, S – susceptible, H – heterozygous.

## Results

The PCR fragments amplified by the primer pairs 0751D06_1 and 0751D06_2 from the parental cultivars of the RIL population (‘CDC Freedom’ (resistant) and ‘Samson’ (susceptible)) were Sanger sequenced from both ends using the same primers as for PCR amplification. The consensus sequences of both fragments were aligned with the full length reference sequence MLOC_38442. Polymorphisms between the sequences from the two parental cultivars were examined, and a SNP in a recognition site for the restriction enzyme PvuII was identified. The recognition site (CAG|CTG) was intact in the resistant parent ‘CDC Freedom’, but in the susceptible parent ‘Samson’ the corresponding sequence was CACCTG, indicating that the PvuII restriction enzyme would not cut the PCR fragment amplified from the susceptible parent. The SNP is located at nucleotide 1592 in the reference sequence MLOC_38442. PCR amplification with the primers flanking the SNP site (see Materials and Methods) resulted in a 170 bp fragment. Digestion of the amplified fragment from the resistant parental variety produced two bands of 111bp and 59 bp, while the analysis of the susceptible parental cultivar produced a single band of 170 bp (PCR product was not cleaved). Heterozygous loci resulted in 3 fragments after PvuII digestion (170 bp, 111 bp and 59 bp).

Subsequently, genotyping was done on the 98 members of the RIL population and 39 RILs had the resistant genotype, 53 the susceptible genotype, and 6 were heterozygotes. The genotyping results were compared with resistance phenotypes from a 2-year artificial inoculation trial using local true loose smut pathotypes. One RIL had a resistant genotype, but was susceptible according to the artificial inoculation results, and of the six RILs with a heterozygous genotype, five were susceptible according to the artificial inoculation results, and one was resistant. The genotyping and artificial inoculation results for the other RILs were consistent.

The newly developed marker was further tested on an additional 47 barley lines and cultivars that were also phenotyped for true loose smut resistance by artificial inoculation. 42 lines and varieties had corresponding genotype and resistance phenotype (15 resistants and 27 susceptible). Of the five lines with inconsistent results, four were susceptible according to the genotyping results, but were resistant according to the artificial inoculation results. This could account for the cases where the genotyping indicated that the line was susceptible, but artificial inoculation was not successful for some reason. One line was resistant according to the genotyping results but had a susceptible resistance phenotype. The analyses for this line should be repeated, to rule out errors, or alternatively, this could represent incomplete segregation between the developed marker and *Un8* true loose smut resistance.

## Discussion

The use of DNA markers linked to genes controlling traits of agronomic interest can increase the efficiency of breeding programs. The development of markers linked to gene alleles conferring resistance to pathogens is of particular utility, as phenotyping for resistance can be difficult, as this necessitates the introduction of pathogens into greenhouses or cultivation areas and artificial inoculation can be inconsistent. In addition, if DNA markers linked to several resistance genes are available, these resistance genes can be pyramided, providing durable resistance to pathogens (Raj and Sanghamitra, 2010). The efficiency of marker assisted selection is determined by the degree of genetic linkage between the marker and the allele of interest. If the markers are not directly within the gene or locus controlling the trait, then genetic recombination between the marker and the gene can lead to inconsistencies between genotype and phenotype. Therefore, markers within genes or loci directly involved in the trait being selected for are desirable, as this minimises the recombination rate, making the marker more consistent within the germplasm where it was developed, and making it more likely to be applicable to a wider range of germplasm.

One important aim of the Latvian barley breeding program is development of varieties suitable for cultivation in organic farming conditions. Therefore, resistance to diseases is essential, since no pesticides are used. Resistance to seed-borne diseases, focusing on loose smut, is one of the main target traits. Germplasm containing *Un8* and other resistance genes is being widely utilised and marker assisted selection can increase breeding efficiency. The first candidate variety containing the *Un8* resistance gene is currently undergoing registration. The marker developed in this study was also utilised for selection of resistant plants in composite cross populations, which were developed using several parents containing the *Un8* resistance gene, with the aim of creating improved heterogeneous populations with resistance to true loose barley smut. Loose smut is also becoming a problem disease in conventional farming, since the seed treatments are losing their efficacy, and the use of pesticides and fungicides is being gradually restricted in the European Union and elsewhere.

A marker for true loose smut resistance was developed based on polymorphism between the barley cultivars ‘CDC Freedom’ (resistant) and ‘Samson’ (susceptible), the parents of a RIL population developed for research purposes. The marker is located in a candidate gene for the *Un8* resistance locus. This gene encodes a tandem kinase protein, but while a marker that is tightly linked to *Un8* co-segregates perfectly with *Un8* in a single mapping population, this candidate gene has not been definitively proved to be the *Un8* resistance gene (Zang et al. 2015; Klymiuk et al. 2021). Tandem kinase proteins have only recently been described as a novel class of resistance proteins (Lu et al., 2020), however, several of this class of genes have been shown to confer resistance to rust (Klymiuk et al., 2018; Chen et al, 2020) and powdery mildew (Lu et al., 2020) in wheat. While further studies are needed to definitively conclude that the candidate gene is involved with true loose smut resistance in barley, the evidence available indicates that this gene is highly likely to be involved in resistance mechanisms.

The genotyping results obtained with the marker developed in this study were mostly consistent with true loose smut resistance determined by artificial inoculation. Only one of the 98 RILs tested had a resistant genotype, but was susceptible according to the artificial inoculation results. Six of the tested RILs were heterozygous for the newly developed marker, suggesting that this locus might be segregating in some of the RILs. Of the six RILs with a heterozygous genotype, five were susceptible according to the artificial inoculation results, and one was resistant. Only three spikes per line were inoculated, and additional tests are needed to determine if resistance to loose smut is also segregating in these lines. Further testing of the marker on an additional 47 lines and cultivars indicated that five lines had inconsistent results between genotyping and artificial inoculation. Four lines were susceptible according to the genotyping results, but were resistant according to the artificial inoculation results and for one line it was the opposite. One source of inconsistencies between genotyping and resistance phenotyping could be infection escapes after inoculation. In the RIL population, one line also had a resistant genotype, but was susceptible according to resistance phenotyping. This could be a result of the marker not segregating with true loose smut resistance, or could be a result of incorrect genotyping or phenotyping. Repeated analysis of inconsistent results is required to confirm or revise these results. Further sequencing of the candidate *Un8* resistance gene from the additional lines and cultivars tested for true loose smut resistance by artificial inoculation will provide additional information about polymorphisms within the candidate gene, and enable comparison with the resistance phenotypes. This, in conjunction with other functional and gene expression studies, will assist in determining if the candidate gene is responsible for conferring resistance to true loose smut resistance.

## Conclusions

The marker developed in this study has been shown to be a reliable indicator of true loose smut resistance in the material tested, the majority of which was germplasm derived from the Latvian organic barley breeding program. Further studies are needed to decisively confirm the role of the candidate *Un8* resistance gene in barley true loose smut resistance, as well as to assess the accuracy of the marker in a wider range on barley germplasm.

## Acknowledgements

This research is funded by the Latvian Council of Science, project “Genetically diverse populations of self-pollinating cereals for organic farming: agronomic performance, effect of environment, and improvement techniques”, project number lzp-2018/1-0404, acronym: FLPP-2018-1.

